# Data-Independent Acquisition Mass Spectrometry to Localize Phosphosites

**DOI:** 10.1101/464545

**Authors:** Qing-Run Li, Shi-Sheng Wang, Hong-Wen Zhu, Fang-Ying Xia, Peter Roepstorff, Jia-Rui Wu, Rong Zeng

## Abstract

In conventional data-dependent acquisition (DDA) mode used for phosphoproteome analysis by mass spectrometry, a considerable number of phosphosites are localized ambiguously. We found that data-independent acquisition (DIA) method could localize phosphosite accurately and confidently due to its acquisition nature of repeated and unbiased mode. Here, we present a robust and promising DIA workflow for modification sites determination. Through dedicated collection and analyses of site-determining ions following their chromatographic profilings, phosphosites could be assigned or rectified unambiguously.

## Introduction

Shotgun proteomics, which largely relies on data dependent acquisition (DDA), have been developed, accepted and widely used to elucidate the intricate nature of proteomes for biological or clinical applications. For DDA method, the parent ions were excluded after several times (usu. one or two) of fragment ions collection (dynamic exclusion) in mass spectrometer. Unluckily, the MS/MS spectra of peptides are usually not acquired when the peptide eluted at its peak in the liquid chromatography run. For unmodified peptides, this type of spectrum is good enough for database searching. Unfortunately, for modified peptide especially phosphopeptides, such an unoptimistic acquisition of MS/MS spectra leads to ambiguous localization of phosphosites, due to lack of enough acquisition of featured fragment ions for phosphorylation location (Figure 1a and 1b)(1). In a typical DDA mode for large-scale phosphoproteome analysis, around 20~30% sites were ambiguously localized (2-4). Therefore, although sophisticated algorithms were developed (5), only limited site-determining ions could be acquired from fewer static spectra in DDA mode. However, it can be seen clearly that, in most prevalent scoring strategies for site localization, the dynamic chromatographic behaviors of phosphopeptides were barely acquired and considered, especially for the featured fragment ions.

**Figure 1.**
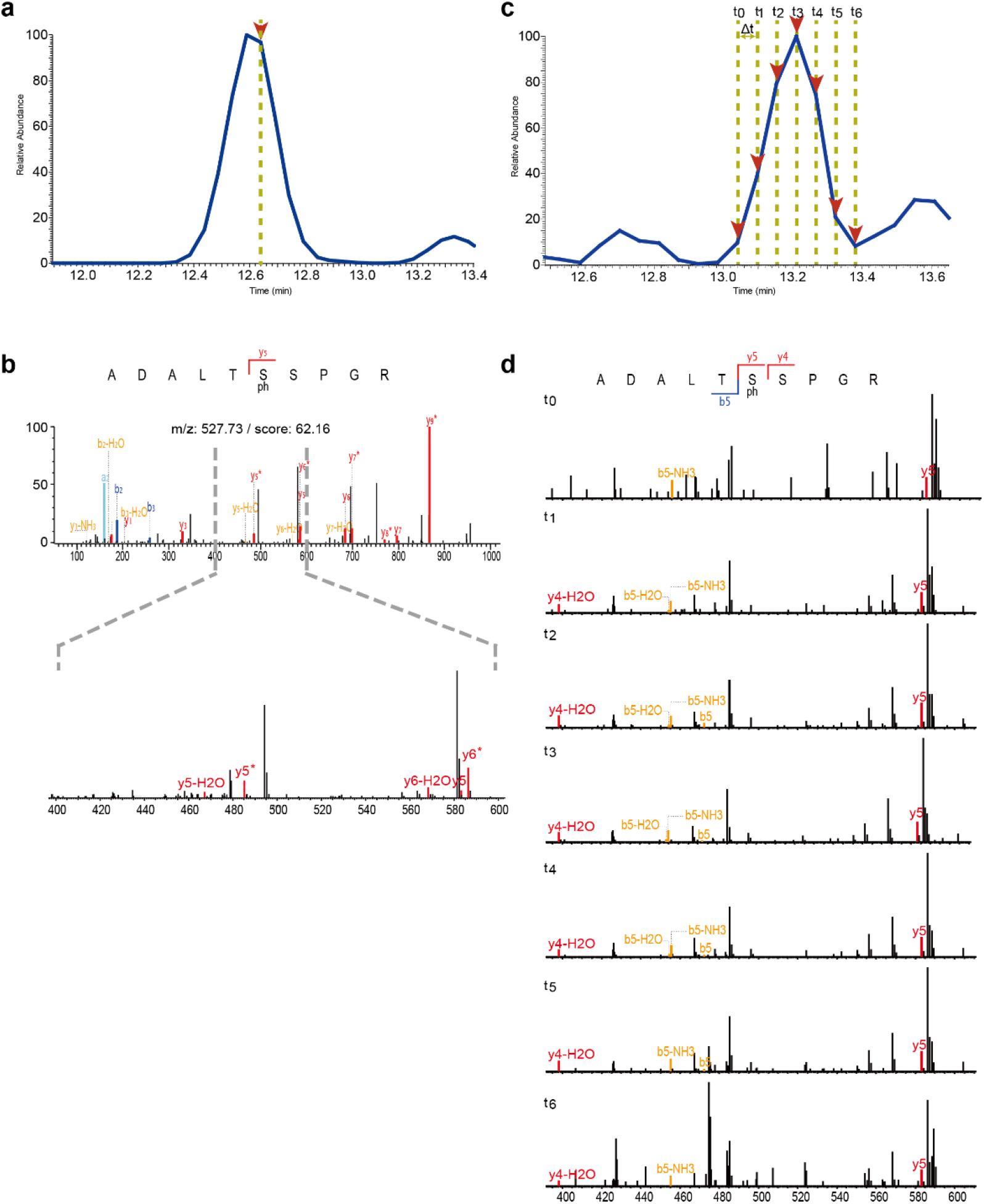
Data acquisition schema for phosphorylation site localization achieved by DDA (left) and DIA (right) methods. For instance, in DDA method, only a few MS/MS spectra were triggered (a) and limited (even missing, in this case) site-determination ions could be obtained (b), which lead to low-confident localization (class II site) for phosphosite. On the contrary, the nature of data collection method allowed as many as important product ions acquired periodically across the LC elution pattern (c). More importantly, in this case, successfully captured the series ions from y5 and y4 could finally localized accurate phosphorylation site correctly (d).

Over the past decade, advances has been achieved in instrumentation technologies for mass spectrometry and chromatography, and the data-independent acquisition (DIA) strategy has been proposed, optimized and applied (6, 7). Although the DIA strategy has been developed and widely used for quantitative proteomics, it has never been reported for correct large-scale phosphosite localization. In DIA approach, the acquisitions of specific isolation window were repeated periodically, which actually allowed for capture of the entire elution pattern not only for one precursor, but significantly for its fragment ions as well.

In the current study, we presented a robust and promising DIA workflow for modification sites determination. Initially, by combining DDA and DIA information simultaneously, we extracted essential features for unbiased phosphosite determination. Through dedicated analyses of site-determining ions following their chromatographic profilings, phosphosites could be finally assigned or rectified unambiguously.

## Experimental Procedures

### Phosphopeptide synthesis

122 phosphosites (Supplemental Table 2) were synthesized (> 95% purity) as internal standards by GL Biochem Ltd (Shanghai, China).

### Generation of phosphopeptides from mouse brown adipocytes

Brown adipocytes were cultured and differentiated as described previously (8). Briefly, the preadipocytes were grown to confluence in culture medium (DMEM + 10% FBS) supplemented with 20 nM insulin and 1 nM T3 (called differentiation medium) (day 0). Adipocyte differentiation was induced by treating confluent cells for 48 h in differentiation medium further supplemented with 0.5 mM isobutylmethylxanthine, 0.5 mM dexamethasone, and 0.125 mM indomethacin. After this induction period (day 2), cells were changed back to differentiation medium, which was then changed every second day till fully differentiated in day 7. Cells were washed thrice with cold PBS buffer and SDS lysis buffer (4% SDS, 100 mM Tris-HCl, 0.1 M DTT, pH 7.6) was used for cell lysis. Cell lysates were broken by sonication, denatured and reduced at 95 °C for 5 minutes. Protein concentration was determined using fluorescence quantification. The proteins were digested by trypsin (Promega) according to the FASP procedure (9). In summary, every 500 μg protein was loaded into a 10 kDa filter, washed twice using 200 μL UA buffer (8 M urea in 0.1 M Tris/HCl pH 8.5) by centrifuging in 12,000 g (the same below), and alkylated using 200 μL 50 mM iodoacetamide (IAA) prepared in UA buffer for 30 min in darkness. IAA was removed by centrifuging and washed thrice using 100 μL UA buffer. Then, the sample was washed twice by 100 mM NH_4_HCO_3_. The final protein pellets were incubated with trypsin (50:1) at 37 °C overnight. Phosphopeptides were enriched with TiO_2_ beads (GL Sciences, Tokyo, Japan) as described previously (10). Briefly, peptides were resolved in 1 mL loading buffer (65% ACN/2% TFA/saturated by glutamic acid) and incubated with TiO_2_ beads for 20 min. A C8-packed tip was used to kept beads and supernatant was removed by centrifugation in 1,500 g. Beads were then washed sequentially with 200 µL wash buffer I (65% ACN/0.5% TFA) twice and buffer II (65% ACN/0.1% TFA) twice. The bound peptides were eluted once with the 200 µL elution buffer I (300 mM NH_4_OH/50% ACN) and twice with 200 µL elution buffer II (500 mM NH_4_OH/60% ACN). The eluates were dried by vacuum freeze, then further analyzed by LC-MS/MS directly.

### LC-MS/MS data acquisition

Both of the synthetic peptides and tryptic phosphorylated peptides were analyzed on the commercial Orbitrap Fusion Tribrid (ThermoFisher Scientific) mass spectrometer, which was coupled to an Easy nLC 1000 system (ThermoFisher Scientific) for sample chromatographic separations, using a binary phase system with phase A that was composed of 2% acetonitrile (ACN), 0.1% formic acid (FA) in water and phase B being 98% acetonitrile, 0.1% formic acid. The peptides separation was performed on a 15 cm column (i.d. 75 µm) packed in-house with the reverse-phase (RP) materials ReproSil-Pur C18-AQ, 3.0 µm resin (Dr. Maisch GmbH, Germany) at 300 nL/min for MS analysis. Peptides were eluted using a linear gradient of increasing mobile phase B (0.1% FA in ACN) from 2 to 35% in 30 minutes for synthetic peptides experiments and 120 minutes for tryptic phosphorylated peptides obtained from mouse brown adipocytes. For DDA, the instrument method consisted of one full MS scan (resolution 240,000 at 200 m/z; 50 ms injection time; 200,000 automated gain control (AGC) target) from 300 to 1500 m/z followed by data-dependent MS/MS scan (resolution 15,000 at 200 m/z; 2 m/z isolation window; 33% normalized collision energy; HCD fragmentation) of a maximum 3 s cycle time with a precursor intensity threshold of 50000. For DIA, the full MS scans were processed in the orbitrap analyzer with the resolution 240,000 at 200 m/z (300 to 1500 m/z; 50 ms injection time; 200,000 AGC target) followed by unequal isolation windows (Supplemental Table S1), which could be calculated by CIW.R (Supplemental Note, Supplemental Figure S1, and Supplemental Figure S2) on the basis of density distribution of precursor m/z in the spectral library (see below), or defined sophisticatedly by the investigator, namely, several MS/MS (resolution 15,000 at 200 m/z; 33% normalized collision energy; HCD fragmentation) after a periodical MS1 scan.

### Spectral library

All DDA data were analyzed with software MaxQuant (1.5.2.8) (11) against a mouse database (2013/07/24) downloaded from Uniprot (http://www.uniprot.org/). For MaxQuant search, the following parameters were used: cysteine carbamidomethylation was selected as a fixed modification, the methionine oxidation, protein N-terminal acetylation, and phosphorylation on serine, threonine and tyrosine were selected as variable modifications. Up to two missing cleavage points were allowed. The precursor ion mass tolerances were 4.5 ppm, and fragment ion mass tolerance was 20 ppm for MS/MS spectra. The criteria used for phosphosite localization were described previously (2), in which a localization score for the phosphosite above or below 0.75 was considered as high (class I) or low (class II) confidence, respectively. The false discovery rate (FDR) of 1% were used for peptide and phosphosite identification. These peptides in msms.txt file were subjected to build a spectral library using Bibliospec, which was integrated into Skyline software.

### Data analysis

All DIA data were firstly analyzed using the open-source software Skyline v3.1.0.7382 to identify a peptide. In the transition settings, charges +2, +3, +4 were matched for precursor ions and +1, +2, +3 for b- and y-type ions of product ions, and ion match tolerance was set to 0.02 m/z. The retention time filtering was conFig.d to use only scans within 5 minutes of MS/MS IDs. In addition, for phosphosites localization in the spectral library, all DIA data spectra information were extracted and preprocessed by an in-house software named “DIADataExtracter”, which was compiled with the C# programming language in the Microsoft Visual Studio 2010 Professional Edition. Two frequently-used preprocessing steps (deisotoping and deconvolution) (12) for high resolution MS/MS data were integrated in the tool. Afterwards, the extracted spectral data were analyzed by the other in-house tool called “DIADataAnalyzer” that was compiled by R language (see Supplementary Notes for details).

## Results

### Rationale

We hypothesized that the information acquired by the DIA would rationally enable accurate site determination for phosphosite with high success rates. Unlike the data recorded in the DDA method (Figure 1a and 1b), in DIA method as demonstrated in Figs. 1c and 1d, the MS/MS spectra of phosphopeptides could be acquired repeatedly. Therefore, this strategy would increase the opportunity for capture the site-determination ions. Simultaneously, incorporated with obtained elution patterns for both parent and product ions, the localization could be more accurately and positively confirmed.

In a proof-of-principle for DIA experiment, the first step was to generate the DDA data to build the spectral library for subsequent ion extraction (Figure 2). We noticed that the valuable DDA data could also be used for peptide m/z density estimation (Figure S1). Given the speed of the current instrumentation, rather than slicing consecutive precursor isolation windows equally from the precursor range (13), we dynamically segmented the DIA acquisition window to take advantage of cycle time efficiently (Table S1). Following the peptide identification by the Skyline (14), we collected only the peptide backbone sequence rather than the modified forms from the static DDA library. Then, the product ions for each possible phosphosite localization were dynamically considered, and scored the related fragmentation ions to extracted the essential features from DIA data (see Figure S8 and Supplementary Notes). Finally, we could achieve the unbiased phosphosite determination combining DDA and DIA information simultaneously and, the FDR was also estimated for correct site localization statistically.

**Figure 2.**
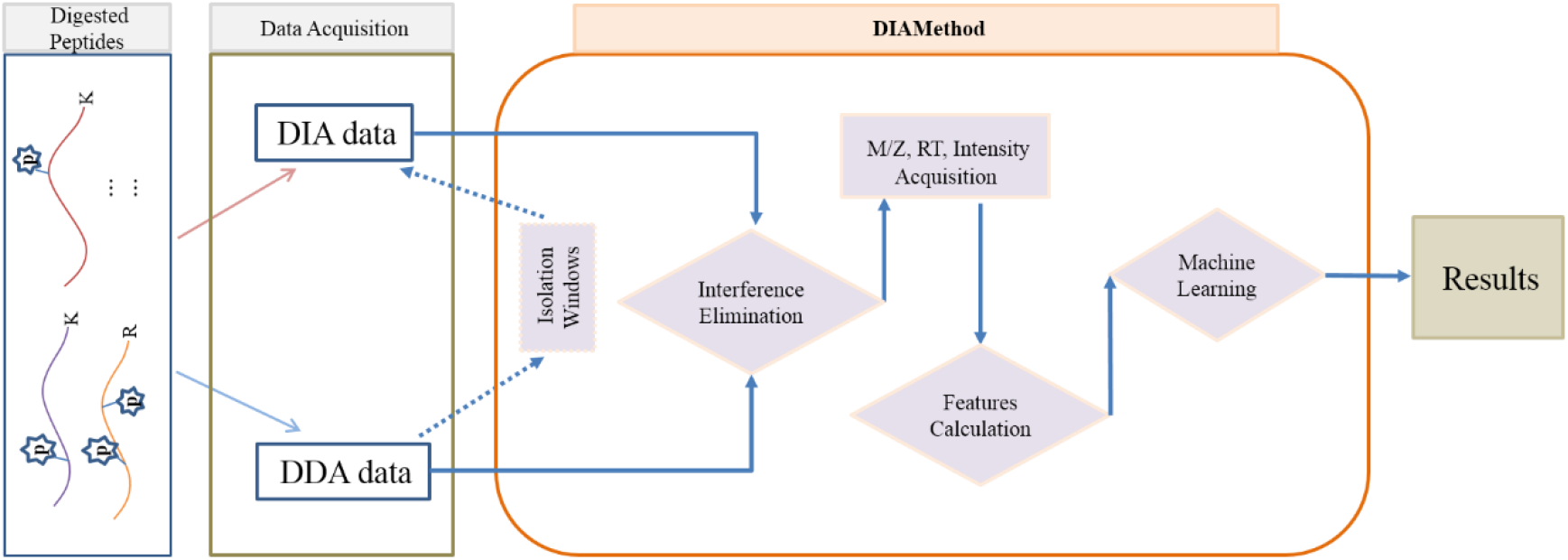
The workflow of DIA method for localization of phosphosites. The DDA data and DIA data were obtained from the digested samples, in which the isolation windows for the acquisition of DIA data could be calculated optionally from DDA data by the module “CIW.R”. Then, the interferences existed in the data could be removed by the tool “DIADataExtracter” and the module “BML.R”. The information of peaks (m/z, retention time, and intensities) could also be extracted by the tool “DIADataExtracter”. Next, the related features were calculated in the modules (“GPPL_ Target.R”, “GPPL_ Normal.R”, “EDIA.R”) and assessed using a supervised machine learning algorithm (support vector machine, SVM) in the module “PHOSITE_ML.R”. Finally, the results were reorganized and output by the module “GPHOSITE_EDIA.R” (see Supplementary Notes for details).

### Localization evaluation by synthesized 122 phosphopeptides

We first assessed the performance of our approach under a simple condition. Altogether, phosphopeptides bearing 122 phosphosites were chemically synthesized to build a reference library (Table S2). Considering the nature of liquid chromatographic and mass spectrometric behaviors for phosphopeptides, three features (Supplementary Notes) were selected, extracted and scored to evaluate the highest-scoring for accurate and sensitive site localization. In each feature, the subscore distributions of phosphosite in target peptides preformed larger values than those in decoy peptides (Figure 3a). Meanwhile, when we combined the three key features together, very low FDR values (0%) were achieved for distinguishing the correct phosphosites from incorrect ones. These results indicated that the high stability and reproducibility of site-determining ions profiles could be detected and parsed by our DIA strategy over the chromatographic profiling (Supplementary Notes). Therefore, as expected, we confidently relocated 108 of 122 phosphosites (Figure S2, Table S3). This result was comparable with DDA method (116 of 122), but fewer false phosphosites identification (6 vs. 29).

**Figure 3.**
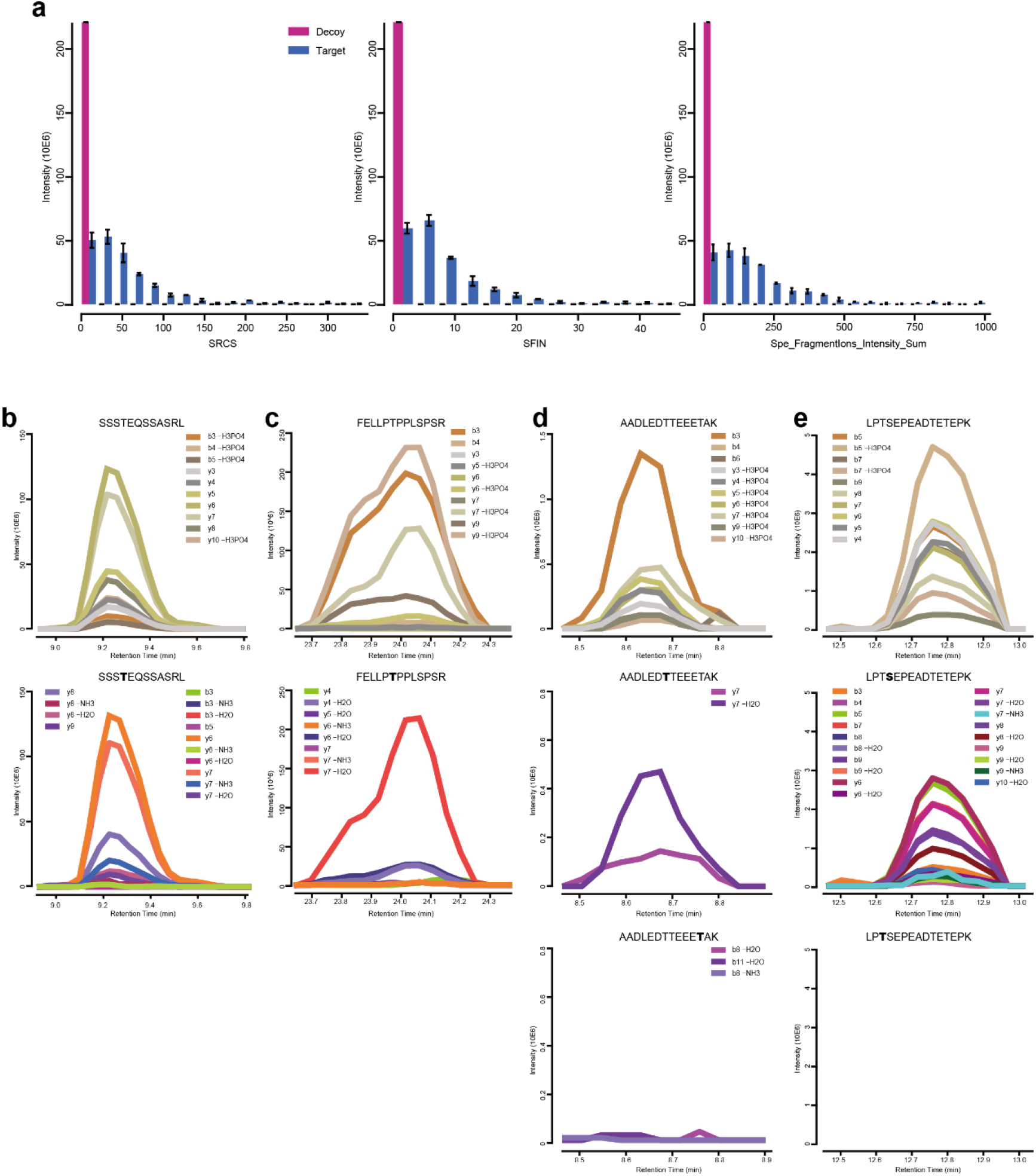
(a) The subscores for three key features distribution of phosphosites in target (blue) and decoy (pink) peptides. (b~e) The graphical demonstration for extracted fragment ion of chromatographic traces: top, the peptide backbone ions; middle, the site-determination ions; down (d and e), the pseudo-decisive fragment ions. Perfect correlation could be profiled between backbone ions (top) and site-determination ions (middle). The bold letter indicates the potential phosphosite.

### Localization confidence evaluation

Next, we examined the confidence of site determination for all identified phosphosite in DIA data. We found the low confident phosphosite (Class II, see Methods for criteria of phosphosite score) identified in DDA method could be verified by our strategy. For peptide SSSTEQSSASRL (Figure 3b, top), the site score for T4 were 0.13~0.55 by multiple identification in DDA, which were classified as low confident phosphosite. However, as fragment ion of chromatographic traces were graphically and clearly demonstrated in our DIA data, the site-determination ions (Figure 3b, middle) were highly correlated with backbone ions (Figure 3b, top), which means high confident and unambiguous phosphosite identification. In this way, up to 93.6% phosphosite with high confident (class I) identification by DDA strategy were confirmed in our DIA results. More importantly, more than 50% lower confident sites (class II) by DDA could also be verified in our DIA strategy when comparing to the site-known reference library (Figure 4a), which further demonstrated the sensitivity and credibility of our strategy.

**Figure 4.**
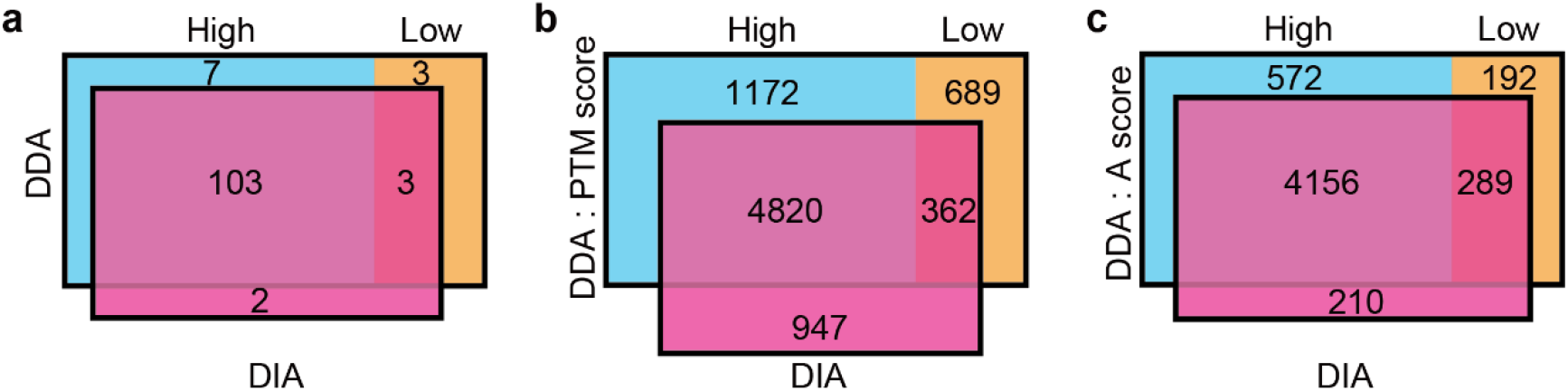
Venn diagram of phosphosites identified by DDA and DIA strategies in (a) synthetic peptides or (b and c) complex samples. Comparison of all phosphosites classified into high confidence and low confidence by DDA method (b: PTM score, c: Ascore) and identified by DIA method from (a) synthetic peptides, (b) mouse brown adipocytes, (c) MCF7 breast cancer cells,(16) respectively.

As we considered all the possibility for potential phosphosites rather than the existing identifications in DDA results, therefore, the missed phosphosites with their site-determination ions in DDA method could be rescued when we checked the synthetic reference library. For instance, T6 phosphorylation in peptide FELLPTPPLSPSR (Figure 3c) was missing in DDA results. However, in DIA method, we could specify the missed localization gracefully with exclusive fragment ions to index the correct phosphosite (Figure 3c, middle).

### Localization incorrect phosphosite

Notably, our workflow could pinpoint the incorrect phosphosite which may be recognized as correct identification in the prevalent DDA method, and *vice versa*. In the DDA-based approaches, various fine-tuned algorithms were developed to exhaustedly localize correct phosphosites from the limited site-diagnostic ions acquired. As shown in Figure 3d, the synthesized peptide AADLEDTTEEETAK, with only one known phosphosite T7, was high-confidently identified in two isoforms with different phosphosites at position T7 (scored 0.78, class I) and T12 (scored 0.99, class I) in DDA method. In contrast, in DIA workflow, because each ion – especially the site-determining ions – could be recorded and analyzed, the correct T7 (Figure 3d, middle) phosphosite could be easily distinguished from incorrect T12 (Figure 3d, down).

### Localization neighboring phosphosite

Intriguingly, as featured site-determining ions from MS/MS could be continuously acquired and extracted following their chromatographic behaviors in our method, we could also successfully distinguish phosphosite for two possible acceptor amino acids which were even placed adjacently(15). The T3 and S4 for phosphopeptide LPTSEPEADTETEPK was identified to be phosphorylated by DDA method with low and high confidence, which were 0.54 (class II) and 0.82 (class I) for localization probability score, respectively. As this peptide was chemical synthesized, we knew the S4 was the only phosphosite in the peptide. In our DIA workflow, all the product ion were extracted, and the probability for all potential phosphorylated amino acid were considered (T3, S4, T10, and T12 in this case). It showed clearly that the decisive fragment ions for localization of S4 (Figure 3e, middle) were perfectly correlated with other backbone b- and y-type ions (Figure 3e, top). While there was no site-determining ion could be extracted to support the localization of the T3 phosphorylation (Figure 3e, down).

### Localization evaluation by large-scale experiments

Lastly, to compare our DIA localization performance to a DDA-based workflow in a large-scale manner, we enriched the mouse adipocyte cell lysate, collected the data from DDA and DIA, and performed the localization analyses. We first examined the key features for precise phosphosite assignment (Figure S3). In accordance with the simple reference library, low FDRs were obtained to distinguish correct from incorrect phosphosites localization. Expectedly, one-third lower confident sites by DDA could be high confidently localized by DIA method (Figure 4b). These results demonstrated a strong and comprehensive separation power to estimate the confidence of phosphosites not only for simple condition, but for complex tryptic peptide mixture as well (Figure S4). Then, from three technical replicates, our DIA assay achieved a deeper overlap between measurements than DDA data (Figure S5), which were 82.8% vs. 74.5%, respectively.

In addition, to avoid possible bias for our workflow to process PTM score for localization, we also analyzed a recently published data set (16), in which another prevalent algorithm “Ascore” (1) was used for phosphosite determination. As expected, more than 80% phosphosites identified with high confidence in DDA were recovered by our DIA workflow. Importantly, 60.1% low confident phosphosites identified by DDA were re-evaluated and verified by DIA with their site-determining ions sophisticatedly aligned following their chromatographic profilings (Fig. 4c, Supplementary Tables 4 and 5). Furthermore, as the site-determining ions achieved from DIA method, we could assigned the confident phosphosite on its peptide backbone, which localizing the phosphoryl group for different phospho-isoforms from DDA data (the same amino acid sequence but different position) (Figure 4). In the DIA method, the intensities of daughter ions could also be extracted for quantification of phosphosites in both synthetic peptides (Figure S6a) and complex samples (Figure S6b). We finally found an improved ability of DIA method for phosphopeptide quantitation reproducibility with a high correlation between measurements (R^2^ above 0.8), and high levels of quantitation accuracy with more than 97% with very low variation (CV below 10%) (Figure S7), when compared with the quantification in DDA strategy.

## Conclusion

In the current study, we took advantage of the nature of the DIA strategy for precise phosphosite localization. After data extraction, the pattern for featured ions of site localization were precisely reconstructed and correlated with the phosphopeptide precursor ion and other ions from the fragment maps. In this way, the dynamic chromatographic profiling was also used for phosphosite determination, rather than limited ions from static spectrum in DDA method. Meanwhile, we are excited to notice that some recent workflow by using DIA data to localize sites for PTMs (17, 18). All of these work proved to be much more sophisticated and subtle for successful phosphosites determination. Our workflow in principle has a potential to be used for accurate localization of other PTMs.

## ACKNOWLEDGMENTS

We thank Dr. Jing Li and Dr. Wei Zhang (Thermo Fisher Scientific, Shanghai, China) for helpful and fruitful discussions. This work was supported by grants from the Strategic CAS Project (XDA12010000), a grant from the Ministry of Science and Technology (2014CB910500) and a grant from National Natural Science Foundation of China (91539124).

## Data Availability

All the raw data of this article are available in the proteomeXchange repository (http://www.proteomexchange.org) with the dataset identifier PXD004368 (Reviewer account details: Username: /Password: irozKOov). The source code for phosphosite localization of DIA data are availability at github.com/wangshisheng/DIADataExtracter_Analyzer.

## Footnotes

The authors declare no competing financial interest.

